# Differentiation Success of Reprogrammed Cells is Heterogenous *In Vivo* and Modulated by Somatic Cell Identity Memory

**DOI:** 10.1101/2024.07.05.602198

**Authors:** Tomas Zikmund, Jonathan Fiorentino, Chris Penfold, Marco Stock, Polina Shpudeiko, Gaurav Agarwal, Larissa Langfeld, Kseniya Petrova, Leonid Peshkin, Stephan Hamperl, Antonio Scialdone, Eva Hoermanseder

**Affiliations:** Institute of Epigenetics and Stem Cells, Helmholtz Zentrum München, German Research Center for Environmental Health, Munich 81377, Germany; Institute of Functional Epigenetics, Helmholtz Zentrum München, German Research Center for Environmental Health, Neuherberg 85764, Germany; Institute of Computational Biology, Helmholtz Zentrum München, German Research Center for Environmental Health, Neuherberg 85764, Germany; TUM School of Life Sciences Weihenstephan, Technical University of Munich, Freising 85354, Germany; Wellcome Trust/Cancer Research, Gurdon Institute, University of Cambridge, UK; Systems Biology, Harvard Medical School, Boston, MA, 02115, USA

## Abstract

Nuclear reprogramming can change cellular fates. Yet, reprogramming efficiency is low and the resulting cell types are often not functional. Here, we used nuclear transfer to eggs to follow single cells during reprogramming *in vivo*. We show that the differentiation success of reprogrammed cells varies across cell types and depends on the expression of genes specific to the previous cellular identity. We find subsets of reprogramming resistant cells that fail to form functional cell types, undergo cell death, or disrupt normal body patterning. Reducing expression levels of genes specific to the cell-type of origin leads to better reprogramming and improved differentiation trajectories. Thus, our work demonstrates that failing to reprogram *in vivo* is cell-type specific and emphasises the necessity of minimising aberrant transcripts of the previous somatic identity for improving reprogramming.

## Introduction

Reprogramming somatic cells into alternative cell fates is a critical aspect of regenerative medicine and stem cell biology. This transformation can be achieved through somatic cell nuclear transfer (NT) to eggs or cloning^1^ and the ectopic expression of specific transcription factors^2,3^. Despite their potential, the clinical use of reprogrammed cells has been hindered by the challenge of achieving an efficient conversion to functional cells fully integrated into tissues^4,5^.

Successful cell fate reprogramming requires erasing the existing cell fate and establishing a new one. On a molecular level, epigenomic and transcriptomic changes are likely key for complete cell fate switches during reprogramming^5^. It has been hypothesised that when the development of NT embryos fails, cell differentiation is defective across cell types due to a failure in epigenome and transcriptome reprogramming of the donor cell fate^6^. Indeed, bulk transcriptome analyses revealed that most NT embryos show an overall persistence of gene expression patterns reminiscent of the somatic donor cell type^7–9^, termed transcriptional memory^6,13^. A distinct set of genes were identified as ON-memory genes, as they were expressed in the donor cell and their expression persisted at unusually high levels in NT embryos when compared to *in vitro* fertilized (IVF) embryos^9^. Additionally, the expected upregulation of many essential genes during embryonic development in IVF embryos was not observed in NT embryos^8,9,11^, which led to their categorization as OFF-memory genes^9^. The retention of high levels of ON- and OFF-memory gene expression at the totipotent stages of NT embryos is indicative of a poor developmental outcome^12,13^. By interfering with chromatin marks associated with reprogramming-resistant ON-memory genes, a reduction of both ON- and OFF-memory gene expression is achieved in NT embryos, correlating with increased developmental success^9,12^. Similarly, reducing chromatin marks linked to OFF-memory genes rescues both OFF- and ON-memory gene expression in cloned embryos and improves their developmental outcome^8,12,14–17^. Collectively, these findings suggest that transcriptional ON- and/or OFF- memory gene expression at the totipotent stage may hamper the successful development of cloned embryos.

Currently, however, the differentiation defects across cell types in developing NT embryos are unknown. Specifically, it is uncertain whether all cell types generated by reprogramming undergo equally defective differentiation or if some cell lineages are more severely affected than others. Furthermore, the role of transcriptional memory in causing these defects, especially in the context of a full organism, is unclear.

In this study, we transplanted endoderm nuclei to enucleated eggs of the frog *Xenopus laevis* to generate NT embryos and monitored the differentiation of the produced reprogrammed cells into epidermal cell types. Our findings demonstrate that the reprogramming outcome and the success of establishing functional cells varies across cell types. While some cell types of the epidermis, such as goblet cells, are formed correctly from reprogrammed cells, other cell lineages, such as basal stem cell-derived ones, show severe differentiation defects. Furthermore, we observed reprogramming resistant cells that retain an endoderm-like state, leading to aberrant body patterning in NT embryos. These phenotypes are accompanied by an increase in cell death. By mimicking transcriptional ON-memory in the epidermis of fertilised embryos, differentiation and body patterning defects analogous to those observed in NT embryos were induced. Conversely, reducing transcriptional ON-memory in NT embryos rescued the observed epidermal defects. These results indicate that transcriptional memory is a key determinant of these phenotypes.

In summary, our study reveals substantial variability in reprogramming efficiency across cell types and identifies the inappropriate expression of lineage-determining genes previously active in the donor nucleus as a crucial obstacle for the generation of functional cell types during tissue development in cloned embryos.

## Results

### Differentiation success varies across epidermal cell types of cloned embryos

Developmental failures in cloned embryos are thought to be caused by widespread cell differentiation defects. We asked whether this is indeed the case, or if specific cell lineages are more susceptible to defects in cloned embryos.

Endoderm nuclei from neurula stage embryos were transplanted to enucleated eggs to generate cloned NT embryos (Fig.1A). Then, cell-fate conversion success in NT embryos was assessed in the developing epidermis. At gastrula stage 12, 2-cell thick layers from IVF and NT embryos were isolated and single-cell RNA sequencing (scRNA-seq) was performed (Fig.1A). Transcriptome profiles from 3,405 cells (1841 IVF and 1564 NT cells) passing quality filters (Fig.S1 A,B) were generated. On average, 42439 unique transcripts and 6329 genes were detected in a typical cell. Unsupervised clustering of the cells’ transcriptomes via the Louvain algorithm identified 10 clusters, which we visualized using Uniform Manifold Approximation and Projection (UMAP)^18^ (Fig.1B Fig.S1C-D). Both NT and IVF embryos contributed cells to all identified clusters (Fig.1C, Fig.S1E). Eight major cell types (Fig.1B) were assigned to those clusters by manually matching the expression of cluster-specific marker genes (Fig.S2A) to *Xenopus tropicalis* scRNAseq datasets^19–21^ or *in situ* hybridization data deposited on Xenbase^22^. Automatic cell type prediction using a *X. tropicalis* single-cell atlas^20^ as a reference confirmed our annotation (see Methods and Fig.S2B). Furthermore, our analysis revealed that both IVF and NT embryos exhibit the greatest transcriptional similarity to X. tropicalis st12 embryos, indicating that any transcriptional differences observed between IVF and NT embryos are unlikely to be attributable to a developmental delay (Fig. S2C). Goblet cells (clusters 1, 2) and cement gland primordium cells (CGP; cluster 4) corresponded to the outer cell layer (*krt* high; Fig.S2D). Cells in the remaining clusters expressed marker genes specific for the sensorial inner layer (*sox11* high; Fig.S2D). There, cell states were identified as non-neural ectoderm (cluster 10), multiciliated cell progenitors (MCPs; cluster 7), basal stem cells (BSCs; clusters 3 and 8) andchordal- and anterior neural plate border cells (CNP and ANP, clusters 6 and 5, respectively; Fig.1B and Fig.S2A).

**Figure 1.**
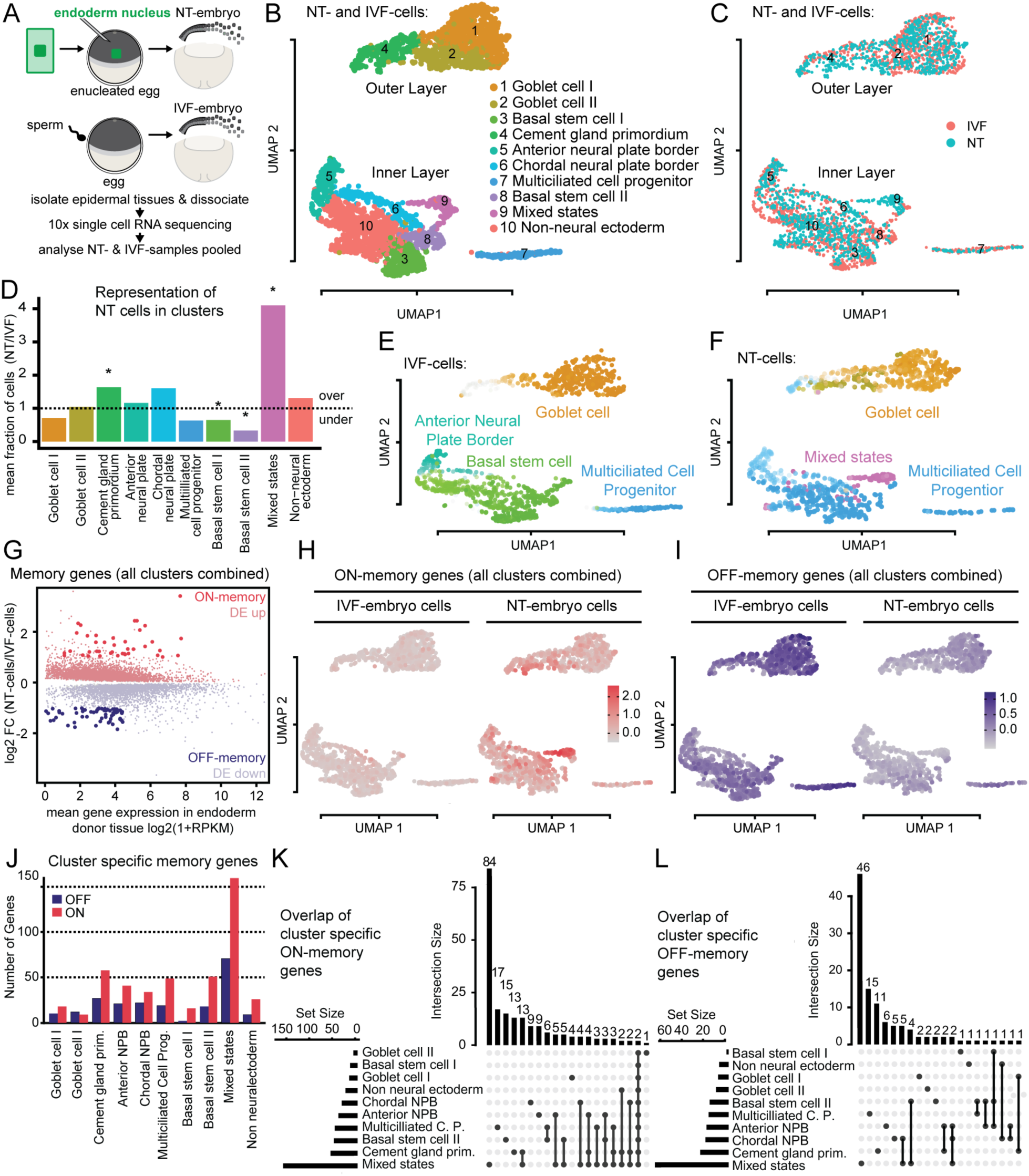
Differentiation defects vary across epidermal cell types and are associated with incomplete transcriptome reprogramming. (A) Design of scRNAseq experiment. (B) UMAP plot of scRNA-seq data from IVF and NT cells coloured by cluster (C) or by condition. (D) Cell type compositional analyses of scRNA-seq data, each bar representing the mean fraction of NT over IVF cells in each cell cluster. over: overrepresented in NT embryos; under: underrepresented in NT embryos. * FDR < 10^-5^. (E) UMAP showing IVF cells coloured by terminal state of differentiation computed using CellRank. Colours match cell clusters shown in B. Darker colours indicate a higher probability of assignment to a terminal state. (F) NT cells analysed and visualized as described in (E). (G) MA plot comparing gene expression between NT and IVF cells. FC: Fold Change; RPKM: Reads Per Kilobase per Million. (H) UMAP plot of ON-memory gene expression in NT and IVF cells. For each cell, the average of the scaled and centred expression levels of the ON-memory genes is shown. (I) Same as (H), showing expression of OFF-memory genes. (J) Bar plot displaying counts of ON- and OFF-memory genes in epidermal cell types of NT embryos. (K) UpSet plot showing overlap of ON-memory genes across cell types in NT embryos. Horizontal bar plot: number of ON-memory genes detected in each cell type. vertical bar plot:number of intersected genes between cell states. Connected dots represent overlap. NPB: neural plate border. (L) UpSet plot showing overlap of OFF-memory genes across cell types in NT embryos as described in (K).

An additional cell cluster (cluster 9) within the inner layer could not unambiguously be attributed to any cell type based on its marker genes (Fig. S2A). Further sub-clustering of this cluster revealed two subpopulations- one with ionocyte characteristics (*foxi1* positive) comprised of both IVF and NT embryo cells, and one with endoderm-like characteristics (*sox17b* positive) consisting mainly of NT embryo cells (Fig.S2 EFG). However, cells of both subclusters express *sox11* (Fig. S2A,D), suggesting that they originate from the inner layer of the developing epidermis. Thus, this cluster 9 is referred to as ‘mixed states’. Overall, our transcriptional analysis suggests that NT embryos can, in principle, form all cell types of the developing epidermis as well as an additional endoderm-like cell type.

Next, we asked if cell states are formed at the same proportions in the developing epidermis of NT when compared to IVF embryos (Fig. 1D). Cell-type composition analyses revealed tht NT cells were significantly overrepresented in cluster 9 with mixed characteristics. Interestingly, NT cells were also significantly underrepresented in the BSC clusters, suggesting that these cell types are formed at a reduced number in NT embryos. Other cell types were represented at comparable rates. Hence, this indicates that not all cell states emerge under similar proportions in the developing epidermis of NT embryos.

The observed changes in cell type composition could be due to differences in the proliferation rate of NT and IVF cells. Therefore, we computationally estimated the cell-cycle phase of the cells within each cluster using our scRNA-seq data^16^. The relative abundance of cells in each cell-cycle phase was comparable between clusters of NT and IVF embryos (Fig.S2H), suggesting that a difference in cell proliferation cannot explain the observed defects in cell-type composition.

To address if differentiation dynamics are altered in NT embryos, and hence could underlie the differences in cell-type composition, we performed single-cell fate mapping using Cellrank^23^. We computed cell-state transition matrices for IVF and NT cells separately and identified terminal states of differentiation. These cell-fate probability maps identified as terminal macrostates, ‘goblet cell’, ‘anterior neural plate border’, ‘basal stem cell’ and ‘multiciliated cell progenitor’ in IVF embryos (Fig.1E, Fig.S2I). In NT and IVF embryos, ‘goblet cell’ was identified within the outer epithelial layer as terminal state (Fig. 1F), suggesting that goblet cells are formed efficiently in both NT and IVF embryos. In the inner layer of NT embryos, we observed the appearance of a new terminal state, ‘mixed states’, and the disappearance of the terminal state ‘basal stem cell’, which agrees with the changes in cell-type composition (Fig.1D). In addition, we also observed in NT embryos a loss of the ‘neural plate border’ terminal state and a shift in differentiation of inner layer cells towards the terminal state ‘multiciliated cell progenitor’. These findings indicate that the differentiation dynamics in NT embryos are disrupted specifically in the cells of the inner cell layer.

In summary, we found evidence that some cell differentiation programs are more vulnerable than others upon reprogramming. Overall, this suggests that the unsuccessful development of NT embryos is linked to a failure in producing specific cell types during development.

### Inefficient transcriptome reprogramming is observed in epidermal cell types with differentiation defects in cloned embryos

Considering that the reprogramming defects occur only in specific cell clusters, we next asked if transcriptional memory of the somatic cell type of origin affects these cell types more than others in the epidermis in NT embryos.

We first characterized transcriptional memory globally in a pseudo bulk differential gene expression analysis. We identified transcripts significantly up- or down-regulated (FDR<0.05) in all NT epidermal cells when compared to all IVF epidermal cells (DE up and DE down; Fig.1G). Then, we identified ON- and OFF-memory genes (see method section) and evaluated their average expression levels across the epidermal cell types. We found that endoderm ON-memory gene expression is highest in cells belonging to cluster 9 with mixed states (Fig.1H), suggesting that this cluster contains cells with high levels of transcriptional memory. Global OFF-memory was instead more evenly distributed across all cell types (Fig.1I). This suggests that transcriptome reprogramming efficiency is not uniform across cell types.

As a complementary strategy to assess differences in transcriptional ON- and OFF-memory across cell types, we determined significantly differentially expressed genes between NT and IVF cells (FDR<0.05) for each cluster separately. From these genes, we identified ON- and OFF-memory genes (see method section) and we found that the “mixed states” cluster 9 possesses the highest number of ON- and OFF-memory genes, corroborating our conclusion that this cluster has relatively high levels of transcriptional memory (Fig.1J). Other cell types with ∼50 or more ON-memory genes include cement gland primordium, multiciliated cell progenitor and basal stem cells. Interestingly, we found evidence of differentiation defects for all these cell types (Fig.1D,E and F). Moreover, we found an enrichment of *X. tropicalis* endoderm markers among the ON-memory genes of the mixed states, cement gland primordium and anterior neural plate border clusters (p-values= 0.03, 0.04 and 0.02, respectively; see Methods), while OFF-memory genes in the multiciliated cell progenitor cluster are significantly enriched for ectoderm markers (p-value= 3 x 10^-5^). Together, this indicates that transcriptional memory varies across cell types and is highest in the “mixed state” cluster 9.

Finally, we evaluated the overlap of ON- and OFF-memory genes across cell types and found that most OFF- and ON-memory genes are cell-type-specific. For instance, 84 ON-memory genes and 46 OFF-memory genes are specific to the mixed states cluster (Fig.1K,L) and another 15 ON- and 3 OFF- memory genes are specific to basal stem cell clusters (FIG.1 K,L). Together, this suggest that each cell type has a specific set of memory genes, and only very few memory genes are shared across multiple cell types.

In summary, we observed cell-type-specific defects in NT embryos associated with high levels of ON- and OFF-memory gene expression, which, in some instances, culminated in cell fate transformations of epidermal cells into an endoderm-like state.

### Endoderm gene expression domains expand into ectoderm regions in cloned embryos

The mixed states cluster 9 contains NT cells that continue to express genes typical of the endoderm donor cell used to generate them (ON-memory genes) as well as genes indicative of an inner layer epidermal fate (ectoderm, *sox11*; Fig.S1H). Such cells with a double identity have not been described so far. Hence, we confirmed their increased occurance in NT-embryos, as suggested by our scRNAseq analyses, and further characterised their localization in intact NT embryos.

We first selected candidate endoderm ON-memory genes as cluster 9 markers. These included the key endoderm transcription factors *sox17b* and *foxa4*, as well as the endoderm genes *march8* and *cdx1* (Fig.2A,D,G and J). We then visualized their expression *via* fluorescence *in situ* hybridization (FISH) or chromogenic whole mount *in situ* hybridization (WISH) assays of endoderm derived NT embryos or IVF-embryos at embryonic stage 12. *In vivo* FISH analyses revealed *sox17b* expression foci in up to 30% of nuclei within the developing epidermis of NT embryos, but not of IVF embryos (Fig.2B and C). WISH against *foxa4* showed that the mRNA was also expressed in cells across the epidermis in NT embryos, but not in IVF embryos (Fig.2E). Furthermore, we could observe that the wild-type expression domain of *foxa4* in the endoderm, which is visible in IVF embryos as a ring around the blastopore, was broadened and extended in most NT embryos until the animal pole and, thus, into the ectoderm (Fig.2E,F). We could also observe the same aberrant and expanded expression domains in NT embryos for *march8* and *cdx1* (Fig.2H,I and Fig.2K,L respectively).

**Figure 2.**
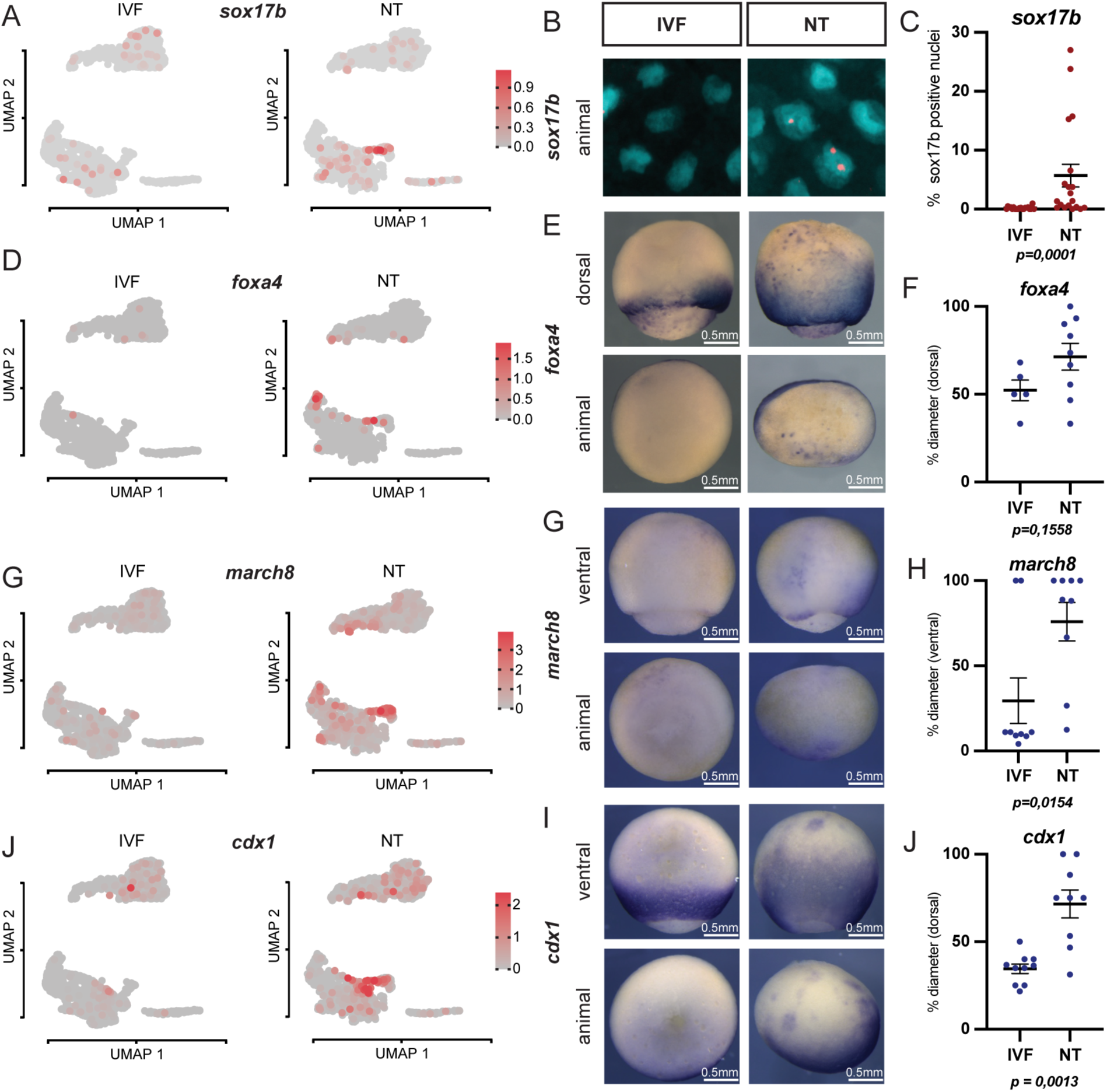
Endoderm gene expression domains expand into ectoderm regions in cloned embryos. (A, D, G, **J)** *sox17b*, *foxa4*, *march8* and *cdx1* expression in UMAP plots. **(B)** Fluorescent *in situ* hybridization against *sox17b* in NT and IVF epithelia; Cyan:nuclei (DAPI); Red:*sox17b* probe (Cy5) **(C)** Quantification of (B) **(E, H, K)** Representative images of IVF and NT st12 embryos stained by whole mount *in situ* hybridization with *foxa4*, *march8* and *cdx1* antisense RNA probes. **(F,I, L)** Quantification of *(E,H,K)*. Per embryo, the ratio of the signal length from dorsal blastopore to animal-vegetal axis diameter was calculated; numbers of embryos for *foxa4* staining: IVF n = 5, NT n = 9; for *march8*: IVF n = 9, NT n = 9; for *cdx1*: IVF n = 10, NT n =9; p-values: unpaired t-test.

Together, these experiments confirm, in NT embryos, the presence of endoderm-like cells in multiple aberrant positions that normally correspond to ectoderm. Notably, the observed expanded expression patterns of these key endoderm genes indicate a disruption of normal embryonic body patterning in NT embryos.

### Basal stem cell numbers are reduced, while epidermal progenitor cells emerge at normal rates in cloned embryos

We then further investigated the defects in epidermal cell type composition, as indicated by our computational analyses of scRNA-seq data, in intact NT embryos.

We observe the expected specification of multiciliated cell progenitors marked by *foxj1* expression (Fig.3A,B), of ionocyte progenitors marked by *foxi1* expression (Fig.3A,C) and of goblet cells marked by *otogl* expression (Fig.3A,D) in our scRNA-seq dataset in IVF and NT embryos. WISH analyses indicate that multiciliated cell progenitors *(foxj1)*, ionocyte progenitors (*foxi1*) and goblet cells (*otogl2*) emerge at similar rates in IVF and in NT gastrula embryos (Fig.3F,H). Basal stem cells (BSCs) are marked by the expression of *tp63*^21,24^ and importantly, their numbers are significantly reduced in the epidermis of NT embryos, when compared to IVF embryos (Fig.3,G,H), confirming our findings of BSC differentiation defects in the scRNA-seq data analyses (Fig.3E).

**Figure 3.**
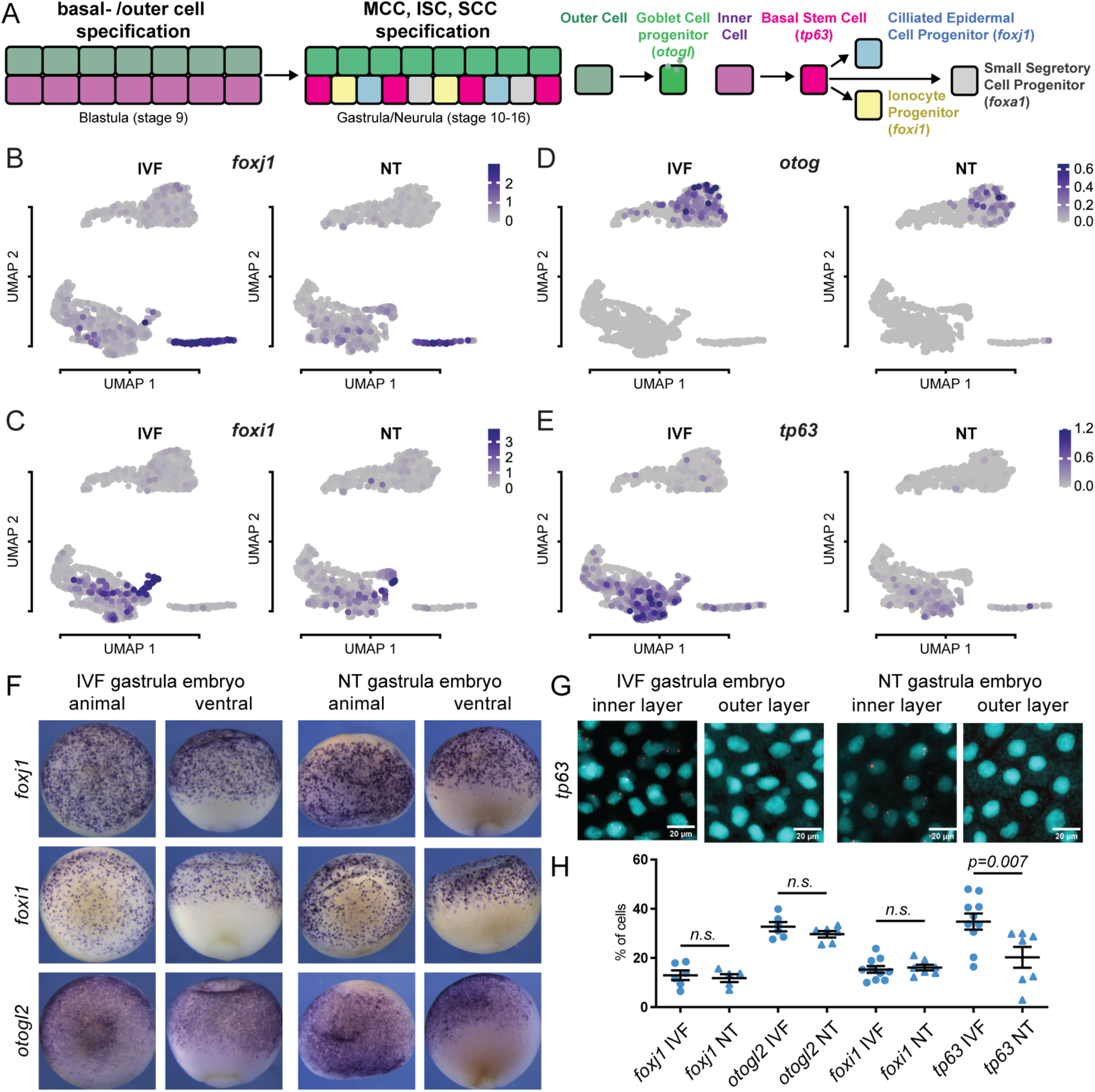
Basal stem cell numbers are reduced in cloned embryos. **(A)** Schematic of cell type specification in mucociliary epidermis **(B-E)** Expression levels of *foxj1*, *foxi1*, *otog* and *tp63* in UMAP plots.**(F)** IVF and NT embryo (st12) whole mount *in situ* hybridization with *foxj1*, *foxi1* and *otogl2* antisense RNA probes.**(G)** IVF and NT epithelia (st12) fluorescent *in situ* hybridization against *tp63* transcript in stage 12 embryos; Cyan – nuclei (DAPI); Red – *sox17b* probe (Cy5); scale bar = 20 µm. **(H)** Quantification of (F,G); *foxj1*: IVF n = 5, NT n = 5; for *otogl2*: IVF n = 8, NT n = 6; for *foxi1*: IVF n = 10 NT n = 7; for *tp63*: IVF n = 10, NT n = 7; p-value: unpaired t-test.

Together, this indicates that during early epidermal differentiation of cloned embryos, the first defect that can be observed *in vivo* is a reduction in BSC numbers.

### Basal stem cell loss and defective mature epidermis coincide with increased cell death in cloned embryos

Basal stem cells could become limiting as growth and differentiation progress and mature epidermal tissues (Fig.4A) are formed. Thus, defects in epidermal cell type differentiation could become more apparent at later developmental stages in cloned embryos.

**Figure 4.**
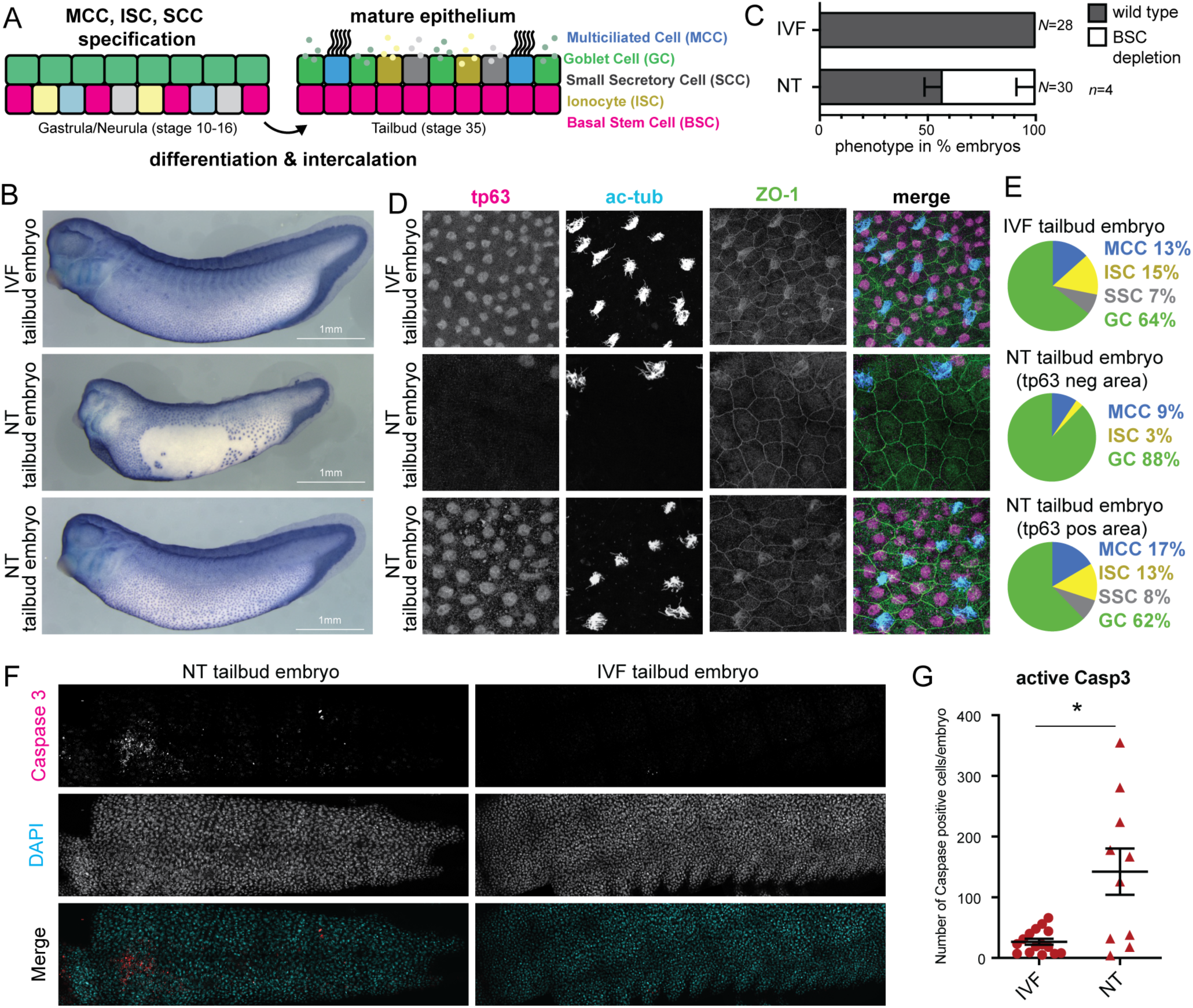
Basal stem cell loss and defective mature epidermis coincide with increased cell death in cloned embryos. **(A)** Schematic of mucociliary epidermis **(B)** IVF and NT embryos stained by whole mount immuno-histochemistry against Tp63; scale bar = 1 mm. **(C)** Proportions of NT and IVF embryos with loss of Tp63 positive basal stem cells (BSC) in epidermis. **(D)** Tp63 positive basal stem cells and α-ac-tubulin positive multiciliated cells in immunofluorescence (IF) staining of epidermis in NT and IVF embryos. Anti -ZO-1 (tight junction protein): cell borders. **(E)** Epidermal cell types in IVF and NT embryos. Data represents mean values from IF stainings in D and data not shown. IVF (n = 7), NT Tp63 negative area (n = 5) and NT Tp63 positive area (n = 5). **(F)** Cleaved Caspase-3 (Asp175) IF stainings of NT and IVF embryos at tailbud stage. Nuclei in cyan (DAPI); cleaved Casp3 in red. **(G)** Quantification of (F). NT (n = 10) and IVF (n = 15).*p < 0.01, unpaired t-test.

To test this, we generated endoderm-derived NT embryos, as well as control IVF embryos, and collected them at the tailbud stage, a time in development when the epidermis has fully matured and differentiated cell types have been formed^25^ (Fig.4A). We first tested if BSC numbers continue to be reduced in NT when compared to IVF using immunohistochemical staining of embryos against the BSC marker Tp63. In 43% of NT embryos (N=30) we observed areas of the epidermis depleted of BSCs, a phenotype never observed in IVF embryos (N=26; Fig.4 B,C). We further observed that BSC derived cell types, multiciliated cells (MCCs), ionocytes (ISCs) and small secretory cells (SCCs), are intercalated between goblet cells in the outer layer of the epidermis at expected frequencies in IVF embryos (Fig.4 D,E, IVF) and in the NT embryos with normal numbers of Tp63 positive BSCs (Fig.4D,E Tp63 positive area). However, in NT embryos with a depletion of BSCs in the inner layer, the BSC-derived cell types MCCs, ISCs and SSCs are missing or reduced in the outer layer and primarily goblet cells can be found (Fig.4D,E Tp63 negative areas). Together, this reveals that NT embryos continue to show BSC reduction in the mature epidermis, which is accompanied by a loss of the mature BSC-derived cell types.

Previously, it has been reported that defective differentiation of BSC-derived cell types results in the depletion of the stem cell pool and increased cell death in the skin of mouse embryos^26^. We next addressed if increased cell death can also be detected in our system by immunostaining against activated Caspase 3. Indeed, most NT embryos showed increased numbers of Caspase 3 positive cells in the epidermis, when compared to IVF embryos (Fig.4F and G). This is indicative of increased cell death in the epidermis of cloned embryos.

Together, this suggests that differentiation of basal stem cells to epidermal cell types is defective in cloned embryos, as we observe a reduction of the stem cell pool, loss of the terminal cell types and increased cell death in the epidermis of cloned embryos.

### Cell differentiation defects are recapitulated by the ectopic expression of ON-memory genes in fertilised embryos

Sox17 and Foxa4 are key endoderm-determining transcription factors^19,27^ that were aberrantly expressed as ON-memory genes in NT epithelial cells. We hypothesised that their expression could contribute to the aberrant basal stem cell differentiation in NT embryos.

Therefore, we tested if overexpression of Sox17b and Foxa4 in the epithelium of embryos generated by fertilisation could phenocopy defects observed in NT embryos. We injected mRNA encoding these transcription factors individually into the ventral blastomeres of 8-cell embryos, which will give rise to the epidermis (Fig.5A). As controls, we injected mRNA encoding a DNA binding protein without known TF activity (Kdm5b^ci^, a catalytically dead histone demethylase) as well as uninjected fertilised embryos. Staining against the BSC marker Tp63 revealed that most embryos expressing Sox17b or Foxa4 in the epidermis showed BSC-depleted regions in the inner layer (Fig.5B,C). As in NT embryos, the outer cell layer located above these depleted areas contained goblet cells but showed a reduction of BSC-derived multiciliated cells, ionocytes and small secretory cells (Fig.5D,E). Furthermore, we found an increase in the number of cells positive for the cell death marker Caspase 3 in the epidermis of embryos ectopically expressing Sox17b and Foxa4, but not in embryos expressing the control protein Kdm5b^ci^ (Fig.5F,G)^ci^.

**Figure 5.**
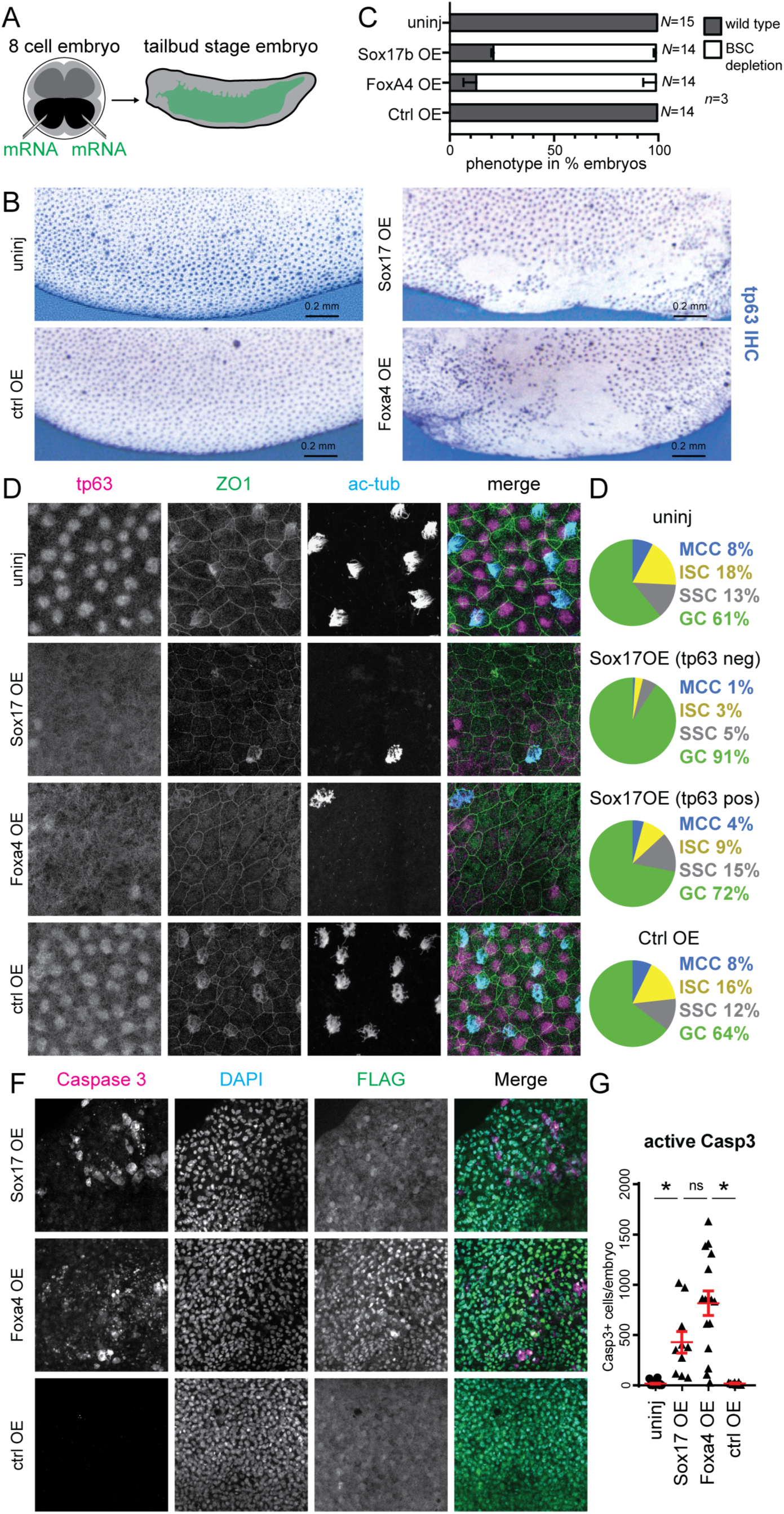
Cell differentiation defects are recapitulated by the ectopic expression of ON- memory genes in fertilized embryos. **(A)** Schematic of microinjection experiment. black: ventral blastomeres. **(B)** Anti-Tp63 immuno-histochemistry of embryos. Uninj.: uninjected, OE: injected embryos overexpressing Sox17b, Foxa4 and Kdm5b^ci^. Scale bar = 0.2 mm. **(C)** Proportions of embryos displaying depletion of Tp63+ basal stem cell (BSC) (white bar) across conditions. **(D)** Immunofluorescence (IF) staining of Tp63+ basal stem cells, α-ac-tubulin+ multiciliated cells and ZO-1+ cell borders in epidermis of uninjected or Sox17b, Foxa4 and control proteins expressing embryos; Tp63: magenta, ZO-1: green, α-ac-tubulin: blue. **(E)** Epidermal cell type quantification in each condition of IF images in (D) and data not shown. Data represents mean values from Uninj, Sox17b OE Tp63 negative areas (n = 5), Sox17b OE Tp63 positive areas (n = 5), ctrl OE (n = 6). **(F)** IF staining for cleaved Caspase-3 and FLAG-Sox17b or FLAG-Foxa4 overexpressed proteins. Caspase 3: magenta, DNA: green, FLAG tagged protein: blue. **(G)** Quantification of (F). Uninj (n = 19), Sox17b OE (n = 10), Foxa4 OE (n = 16), ctrl OE (n = 7).).*p < 0.01 unpaired t-test.

Together, these data suggest that the ectopic expression of endoderm ON-memory genes can induce the same epithelial defects we identified in NT embryos.

### Reducing expression of the key endoderm ON-memory gene Sox17b rescues epidermal defects in endoderm derived NT embryos

Finally, we tested whether reducing the aberrant expression of Sox17b in epithelial cells of NT embryos is able to rescue the observed epithelial phenotypes in NT embryos.

We inhibited Sox17b translation in the developing epidermis by injecting *sox17b* antisense morpholinos (asMOs) into the ventral blastomeres of 8-cell embryos. Embryos were generated by NT of endoderm cells to enucleated eggs or by *in vitro* fertilization (Fig.6A). To trace asMO targeted areas in NT embryos, we co-injected fluorescently labelled dextran. We collected embryos at the tailbud stage and identified BSCs using Tp63 immunostainings. In NT embryos with successful targeting of *sox17b* asMOs to the epidermis, as indicated by dextran-positive cells, we observed a wild-type representation of epidermal BSCs in the epidermis in 8 out of 9 embryos (n=3; Fig.6BC) indistinguishable from IVF embryos (N=15 embryos, n=3; Fig.6BC). Instead, in control MO injected NT embryos, BSC-depleted regions of the epidermis were observed in 4 out of 9 embryos (n=3; Fig.6BC). This suggests that reducing Sox17b ON-memory gene expression in the epidermis of endoderm derived NT embryos restores BSC numbers.

**Figure 6.**
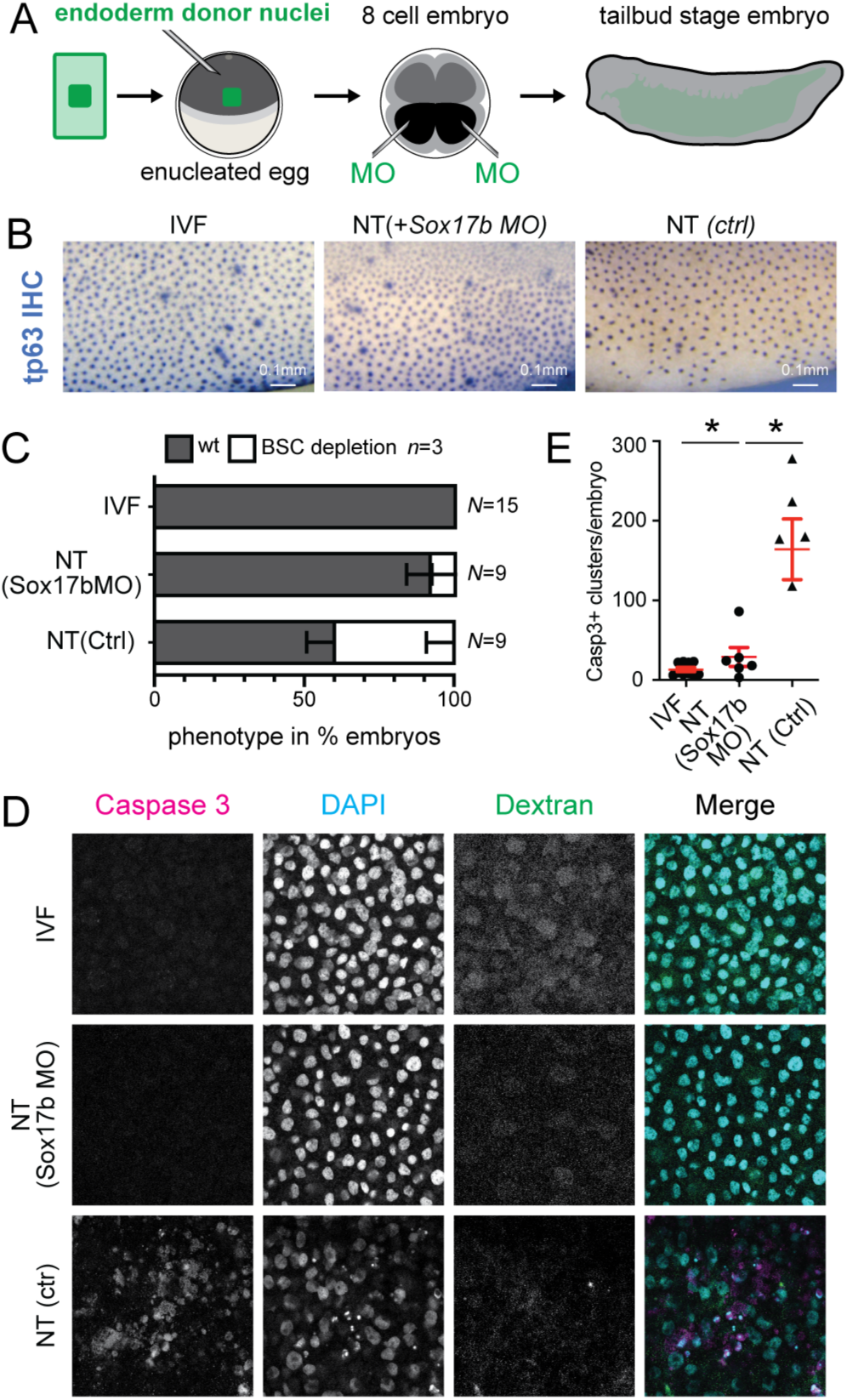
Reducing expression of key endoderm ON-memory gene Sox17b rescues epidermal defects observed in endoderm derived NT embryos. **(A)** Schematic of rescue experiment. NT embryos were injected with antisense morpholinos (MO) into ventral blastomeres (in black) at 8-cell stage. **(B)** Immunohistochemistry for Tp63 protein at tailbud stage embryos. Scale bar = 0.1 mm. Ctrl - control, MO - antisense morpholino. **(C)** Proportions of embryos showing perturbations in the composition of Tp63+ cells in the epidermis. **(D)** Immunofluorescence for activated Caspase-3 (Magenta), DAPI (Cyan) and fluorescent dextran (Green) in IVF and NT embryos injected with control or *sox17b* morpholino. **(E)** Quantification of (D).*p < 0.01 unpaired t-test.

Next, we examined the number of apoptotic cells in these embryos and found that inhibiting *sox17b* ON-memory gene expression in the epidermis by asMO injection reduced the number of caspase-positive cells per embryo to levels similar to those observed in IVF embryos (n=2; Fig 6D,E). Instead, 5 out of 6 untreated NT embryos (n=2; Fig.6D,E) showed elevated numbers of apoptotic cells, as observed before (see Fig.4F,G).

Together, the data suggest that a reduction in the expression of the endoderm ON-memory gene Sox17b rescues epidermal defects in endoderm-derived NT embryos. This, in turn, indicates that ON-memory gene expression is a major contributor to the abnormalities observed in NT embryos.

## Discussion

Our study reveals that reprogramming success *in vivo* is cell-type-specific and uncovers memory of active transcriptional states as both an indicator and an underlying cause for the observed developmental problems in NT embryos.

By leveraging NT in the frog model system, we followed the development of reprogrammed cells within an organismal context and described their *in vivo* differentiation pathways. We uncover a previously unappreciated heterogeneity of differentiation success across cell-types in NT embryos. Being able to compare the single-cell transcriptomes of reprogrammed cells in NT embryos to their exact *in vivo* counterparts allowed in-depth computational analyses of the data generated here, including determining the frequency of NT and IVF cells within the cell clusters and cell differentiation dynamics. These analyses reveal that, surprisingly, many cell states differentiated normally in NT embryo. However, the relative abundance of basal stem cells was reduced in NT compared to IVF embryos, and their ability to differentiate was disrupted in our *in silico* analyses. Moreover, we found a new cell state formed exclusively in NT embryos, which co-expressed epidermis and endoderm markers and altered the tissue’s differentiation dynamics. Together, these analyses singled out differentiation defects in specific types of reprogrammed cells, which we confirmed *in vivo* in NT embryos. This suggests that embryonic death observed in cloned organisms is not due to an overall failure in differentiation of all cell types, but rather the result of defective differentiation of specific cell lineages only.

The single-cell resolution of our study led to the discovery that ON-memory, as well as differentiation defects, are not equally present in every cell of an NT embryo, but are especially prominent in specific cell types, e.g., in the cells with a mixed state and in basal stem cells. We speculated that NT cells with mixed endoderm/ectoderm identity in an epidermal environment could result in aberrant body patterning of the developing embryos. The observed expansion of endoderm gene expression domains far into normally ectoderm regions confirmed this *in vivo*. Furthermore, we hypothesized that basal stem cell defects in NT- embryos might be driven by the high degree of transcriptional memory. Indeed, reducing the expression level of one key lineage determining transcription factor showing ON-memory, Sox17b, increased the differentiation success of reprogrammed cells. We propose a functional hierarchy amongst the genes showing transcriptional memory in cells of NT embryos: We previously found that sox17b is a gene especially resistant to reprogramming due to the stabilization of its active epigenetic state by chromatin modifications such as H3K4me3^9^. When erroneously expressed in epidermal cells of NT embryos, due to epigenetic ON-memory, we speculate that Sox17b induces expression of other endodermal genes and disrupts the normal activation of epidermal gene expression networks, such as the one of basal stem cells and their progeny, thereby preventing normal differentiation. Other cell lineages, such as goblet cells, may more effectively reprogram endodermal cell fate into their own lineage. However, the factors that determine why certain differentiation programs in reprogrammed cells are able to overcome the epigenetic memory of the donor cell type in NT embryos remain unclear.

We observed that NT embryos also show an increase in caspase-positive cells, suggesting an increase in cell death. Our discovery that the expression levels of Sox17b in epidermal tissues of IVF and NT embryos correlate with cell death indicates a link between transcriptional ON-memory and cell death. Interestingly, in zebrafish, cells with a transcriptional signature that does not fit the surrounding tissue become apoptotic via cell competition^28^. Phenotypes similar to those we observed in frog NT embryos are also present in the developing epidermis of mouse embryos when cell competition is induced^26^. Hence, it is tempting to speculate that the observed poorly reprogrammed cells with high degrees of ON memory are eliminated by the more efficiently reprogrammed cells in NT embryos through cell competition.

Overall, our results provide new insights into the outcomes of cellular reprogramming *in vivo* and its molecular drivers, which are crucial for regenerative medicine. Reducing the expression of specific ON-memory genes emerges as a novel strategy for efficiently producing functional cells and tissues capable of replacing irreversibly damaged tissues through cell reprogramming.

## Experimental Procedures

### Resource Availablility

Corresponding authors: E.H.,A.S. Materials availability: Produced Materials are available upon request. Data and code availability: The data have been deposited at GEO (GSE269252). Original code can be accessed at https://github.com/ScialdoneLab/scXen.

### Xenopus Laevis

*Xenopus Laevis* were obtained from Xenopus1 (Xenopus 1, Corp. 5654 Merkel Rd. Dexter, MI. 48130). Frog care was conducted according to German Animal Welfare Act and in accordance with guidelines approved and licensed by ROB-55.2-2532.Vet_02-23-126.

### Embryo handling and nuclear transfer

*Xenopus* eggs were collected*, in vitro* fertilized and handled as described^29^. Developmental stage of embryos was determined according to Nieuwkoop and Faber^30^. Donor cells were isolated from endoderm tissue of neurula stage embryos (stage 21) and transplanted to enucleated eggs as describen in Hoermanseder et al.^9^

### Single cell preparation

Epidermal tissues were isolated from st12 IVF and NT embryos, washed and resuspended in Newport 2.0 dissociation buffer (100mM sodium isethionate, 20mM sodium pyrophosphate, 10mM CAPS, 20mM glucose, pH10.5NaOH), transferred to BSA coated microcentrifuge tubes and incubated for 30min at 18°C with agitation. Single cell suspensions were resuspended in PBS-BSA and filtered through 30 µm cell strainers. Cells were washed 1 ml PBS-BSA, counted and analysed for viability. 2500 cells with viability more than 95% and not detectable RNA in supernatant were used for library preparation.

### Single cell capturing, barcoding and library preparation

In 2 separate experiments, single cell suspensions from pools of 5 IVF or 5 NT epidermal tissue samples were processed using Chromium 10x Genomics platform to generate single cell libraries. Libraries from all samples were pooled and sequenced in two lanes using Illumina HiSeq 2500 to generate paired-end 100-bp data.

### scRNAseq data pre-processing

Single cell libraries were processed using the 10x Genomics Cell Ranger pipeline (v2.2.0 and v3.0.2 for 10x chemistry 2 and 3 respectively) and aligned to *Xenopus laevis* genome (v9.1) with STAR^31^. Number of cells was estimated from the distribution of barcode counts and data realigned using --force-cells option. Quality control was based on the mean number of detected genes (3k-6k) and mean UMIs per cell (80k-500k). Mapping efficiency was high, with >75% reads mapped to the genome, and 40-60% mapped to the transcriptome. After quality control, 3405 cells (>6k UMIs and >2k genes per cell) were retained: 566 for SIGAA2 (IVF1), 514 for SIGAB2 (NT1), 1275 for SIGAH5 (IVF2) and 1050 for SIGAH12 (NT2), for a total of 1841 IVF and 1564 NT good quality cells.

### Data normalization, identification of highly variable genes and batch integration

Data for each batch was normalized with “NormalizeData” from the Seurat v3^32^ package, and the top 2000 highly variable genes were computed (“FindVariableGenes” function, “vst” method). The data for the four batches were integrated with “FindIntegrationAnchors”, using the first 20 dimensions from the Canonical Correlation Analysis (CCA), and “IntegrateData”. The integrated data was scaled with “ScaleData”.

### Dimensionality reduction and clustering of cells

Principal Component Analyses (PCA) and a UMAP (function “RunUMAP”) were computed on the integrated and scaled data. A k-nearest neighbour (KNN) graph (function “FindNeighbors”) was computed and cell clustering was performed (function “FindClusters”). Number of neighbors and resolution were chosen using information theoretic criterion defined in^33^ and “resolution”=0.5 and “k.param=20” (Fig.S1C) was selected, which identified 11 cell clusters. Cells from cluster 0 were excluded due to low UMI and gene counts (Fig. S1A-B), as were14 outlier cells from cluster 6 identified with “PyOD”^34^. Cells from cluster 9 were subclustered with the batch integration pipeline from Seurat v3 applied as described above and cell clustering with default parameters was performed, obtaining two clusters.

### Cell type annotation

Cell clusters were annotated based on the expression of well characterised eidermal marker genes of Xenopus cell types at stage 12 using the *Xenopus* Bioinformatics Database (Xenbase)^22^ and *Xenopus* datasets ^19–21^. For automatic cell type annotation, *Xenopus tropicalis* single-cell atlas^20^ (GSE113074) was used as reference. *X. laevis* gene names were mapped to *X. tropicalis* using a pipeline of reciprocal gene symbol comparison^35^. Expression of genes from the long (.L) and short (.S) chromosomes was aggregated by taking the mean. Cells were mapped onto the *X. tropicalis* atlas at stage 12 (reference data) using Scibet R^36^. Predicted cell types for each cluster from the first analysis step were considered and reference clusters of the same cell type at various developmental stages from the *X. tropicalis* atlas were gathered. Our clusters were projected onto this reference framework via Scibet, separately for IVF and NT embryos, to predict the closest corresponding developmental stage for each cluster. For clusters 2,4,7 and 9, we did stage mapping using the top two predicted *X. tropicalis* cell types since they had similarly high scores (Fig.S2B).

### Cell cycle annotation

The cell cycle phase was annotated using the “cyclone” function (“scran” R package^37^).

### *In silico* cell type composition analysis

Changes in cell type proportions between IVF and NT were tested following the method of ^38^.

### CellRank analyses

Fate mapping analysis on the scRNA-seq data was conducted with CellRank (v1.1.0)^23^. Since there is not a well-established method to take into account batch effects in RNA velocity analysis, only the SIGAH5 (IVF) and the SIGAH12 (NT) samples were considered due to their larger number of cells. Matrices of spliced and unspliced counts were computed using velocyto^39^, “scvelo” ^40^ was used to compute RNA velocities using default parameters and dynamical model. RNA velocity confidence was estimated using “scvelo.tl.velocity_confidence” and represented using “boxplot” function of “seaborn” package. Confidence scores were high for all clusters (Fig. S1M), which supported the use of RNA velocity in CellRank, as described below. We used the CellRank function “cellrank.tl.terminal_states” (with default parameters) to infer terminal states of cell dynamics based on velocity and the connectivity kernels. Plots of terminal states for IVF and NT cells were obtained using “cellrank.pl.terminal_states” function, where darker colors correspond to higher-confidence estimations.

### ON and OFF memory genes identification and data representation

Differential expression analysis was performed between IVF and NT cells, across all clusters combined, using the R package “DESeq2”, while controlling for the experiment (SIGAA2-SIGAB2 vs SIGAH5-

SIGAH12) and with a significance threshold FDR<0.05. Then, relying on the bulk RNA-seq data from^9^, we defined global ON- and OFF- memory genes using the following criteria: ON-memory genes: mean(RPKM_Donor_)> 1 (from bulk RNA-seq); FDR_Donor vs IVF_ < 0.05 (from bulk RNA-seq); log2(FC)_NT/IVF_ > 1 (from scRNA-seq); OFF-memory genes: mean(RPKM_Donor_)< 20 (from bulk RNA-seq); FDR_Donor vs IVF_ < 0.05 (from bulk RNA-seq); log2(FC)_NT/IVF_ < -1 (from scRNA-seq). The same analysis was performed considering each cell cluster separately, and cluster-specific ON- and OFF- memory genes were defined using the same criteria above. To characterize the sets of cluster-specific memory genes, a list of cell-type markers from the *X. tropicalis* single-cell atlas^20^ was used. Lists of markers at the germ layer level (endoderm, ectoderm, mesoderm) were aggregated, cluster-specific ON- and OFF- memory genes were considered and *X. laevis* gene names were mapped to *X. tropicalis*, as described above. For each cluster-specific ON- and OFF-memory gene list, the enrichment for endoderm, ectoderm or mesoderm markers was tested with a Fisher’s exact test, using as a background the set of genes tested for differential expression. P-values for multiple testing were adjusted with Benjamini-Hochberg correction.

### Whole mount RNA *in situ* hybridization

**F**or RNA *in situ* hybridization analysis, embryos were anesthetized in 0.05% Tricaine, fixed 1h at RT in MEMFA (0.1M MOPS pH7.4, 2mM EGTA, 1mM MgSO4, 3.7% Formaldehyde), washed with PBS and stored in ethanol at -20°C. Primers used for amplification of selected genes are listed in Table S1“List of used primers”. All amplicons were subcloned into pCS2+ vector, linearized, transcribed using RiboMAX™ kit (Promega, #P1300 and purified (RNeasy Mini-Kit; Qiagen, #74106). Fluorescent RNA *in situ* hybridization used tyramide amplification after addition of probes and incubation with horseradish peroxidase antibody conjugate (Sigma-Aldrich, #11207733910) as described in Lea et al. (2012)^43^. Chromogenic RNA *in situ* hybridization was performed as described in Sive et al. (2000)^32^. After the AP staining (BM purple, Sigma-Aldrich, #11442074001) the embryos were dehydrated with 75%EtOH/PBS and bleached 1-2h in bleaching solution (1% H2O2, 5% Formamide, 0.5xSSC). After refixation in MEMFA, the embryos were photographed using a Leica M205FA stereomicroscope. All images are presented as a compound z-stack projection.

### Immunohistochemistry for Tp63 protein

Embryos at stage 32-33 were fixed overnight at 4°C in MEMFA, rinsed with 1xPBS and incubated at 24 hrs in 100% ethanol at -20°C. Embryos were then rehydrated with stepwise washes using 75%, 50% and 25% ethanol in PBST (1xPBS+0.1% Tween20). Endogenous AP was inactivated by washing embryos once with 50% formamide/PBS solution and incubating them in the same solution at 65°C for 2 hrs. Embryos were then permeabilized twice 10 min in 1xPBS+0.2% TritonX-100, blocked in antibody buffer (1%BSA in 1xPBS+0.02% Tween20) for 1 h at RT and stained overnight (o/n) with primary antibody anti-Tp63 (Abcam, clone 4A4, #ab735) at 4°C, followed by PBST washes (6x1h), 1 h of blocking with antibody buffer and o/n incubation with anti-mouse AP conjugated antibody (Abcam, #ab97262) at 4°C, PBST washes (6x 1h), incubated 20 min in AP buffer (100mM Tris/HCl pH=9.5, 100mM NaCl, 50mM MgCl2) and stained with NBT (33.8 µg/ml)/BCIP(17.5 µg/ml) in AP buffer for 1.5 h at 4°C. After the signal was developed, embryos were fixed 30 min in 4% PFA/in 1xPBS solution, dehydrated with 30 min with 75% ethanol in 1xPBS and bleached for 2-4 h until Tp63 positive cells were visible.

### Immunofluorescence

Detection of epidermal cells was performed as described in Walentek et al. (2018)^25^. Shown IFs are: Anti-Acetylated Tubulin (Sigma-Aldrich, #T6793); anti ZO-1 (Invitrogen, #61-7300), β-catenin (rat, hybridoma supernatant gifted from prof. Ralph Rupp); anti-cleaved Caspase-3 (Asp175) (CST, #9661), anti-FLAG (mouse, Sigma-Aldrich, #F3165), anti-mouse-AF488, anti-rabbit-AF555 or 647, anti-rat-AF546, all from Thermo-Fisher Scientific, DAPI and PNA-lectin AF594 (Invitrogen, #L32459). Before mounting into Vectashield (Vector labs, H-1000), embryos were washed 3x30 min in TBST. Imaging was conducted on Leica SP8 system using oil 40x objective, LASX software, z-step 0.7µm, 1024x1024 per image, 0.75x zoom.

### mRNAs and Morpholinos

mRNA for *sox17b.1.S*, *foxa4.L* and mouse *Kdm5b^ci^* transcription factors was generated from pCS2+ vectors, where coding sequence of all genes was fused with *Xenopus* globin 5’ and 3’ UTRs and tagged NT 3xFLAG. RiboMAX™ Large Scale SP6 RNA polymerase kit (Promega, #P1280) was used for *in vitro* transcription on XbaI linearized pCS2+ vectors. 250 pg of mRNA in 4.6nl was injected with 100 pg of fluorescent dextran AF488 (Invitrogen, #D22910) into both ventral blastomeres of 8-cell stage embryos. Morpholino oligo *sox17β* described in Clements et al. (2003)^47^ was injected in concentration of 5 ng per 8-cell stage ventral blastomere of NT or IVF embryos.

**Table 1.**
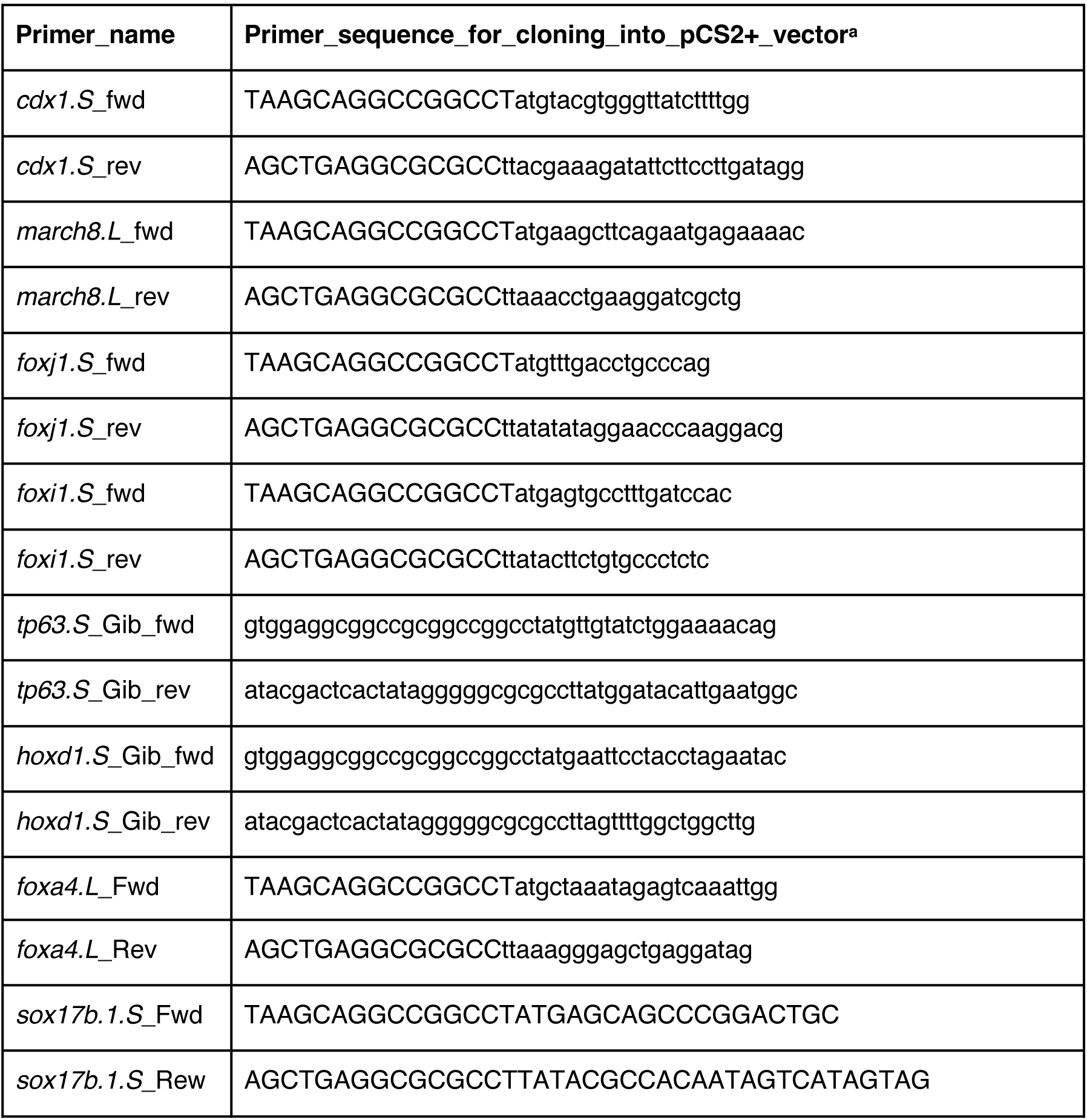
List of used primers.

## Acknowledgements

T.Z. and E.H. were supported by CRC1064 and HO 6864/2-1, both DFG; Project grant MR/P00479/, MRC. S.H. by ERC starting grant 852798. Work in the labs of E.H., A.S. and S.H. was funded by the Helmholtz Association. J.F. and M.S. were supported by the Joachim Herz Stiftung Add-on Fellowship for Interdisciplinary Life Science. M.S. was supported by “MUDS”. P.S. was supported by the Helmholtz Munich epigenetics summer internship program. We acknowledge data generated in XENCAT atlas OD031956. We thank J.Gurdon, J.Jullien, C.Bradford, R.Rupp and his team, P.Walentek, M-E. Torres-Padilla, A.Burton, M.Oak and members of our team and institutes for reagents, input and support.

## Author contributions

T.Z., J.F., C.A.P, G.A., P.S. and E.H. conducted the experiments and analyses; K.P and L.P. bioinformatics support; S.H. provided financial support; T.Z., J.F., E.H. and A.S. designed the analyses and experiments; E.H, A.S, J.F. and T.Z. wrote the paper.

---

**Supplementary figure 1.**
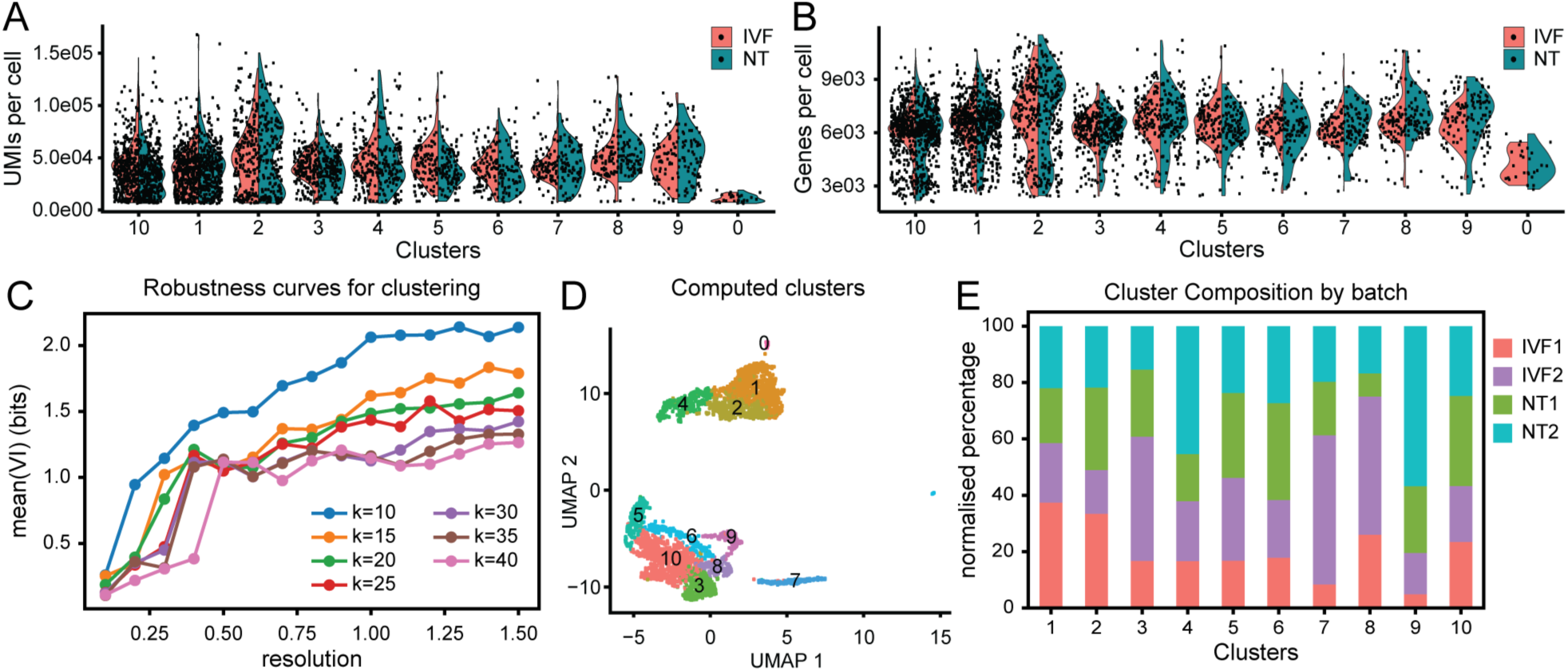
Differentiation defects vary across epidermal cell types and are associated with incomplete transcriptome reprogramming. **(A)** Number of unique molecular identifiers (UMls) identified per cell for each cluster. Coloured by condition IVF and NT. **(B)** Number of genes identified per cell for each cluster. Coloured by condition IVF and **NT. (C)** Variation of Information (VI) between cell clusterings, averaged over M=SO random sub-samplings of the highly variable genes, versus the resolution parameter of the Louvain algorithm. Colours indicate different values of the number of neighbours used to compute the Shared Nearest Neighbour (SNN) graph of the cells. **(D)** UMAP plot of the scRNA-seq data coloured by cell clusters identified via Louvain algorithm.

**Supplementary figure 2.**
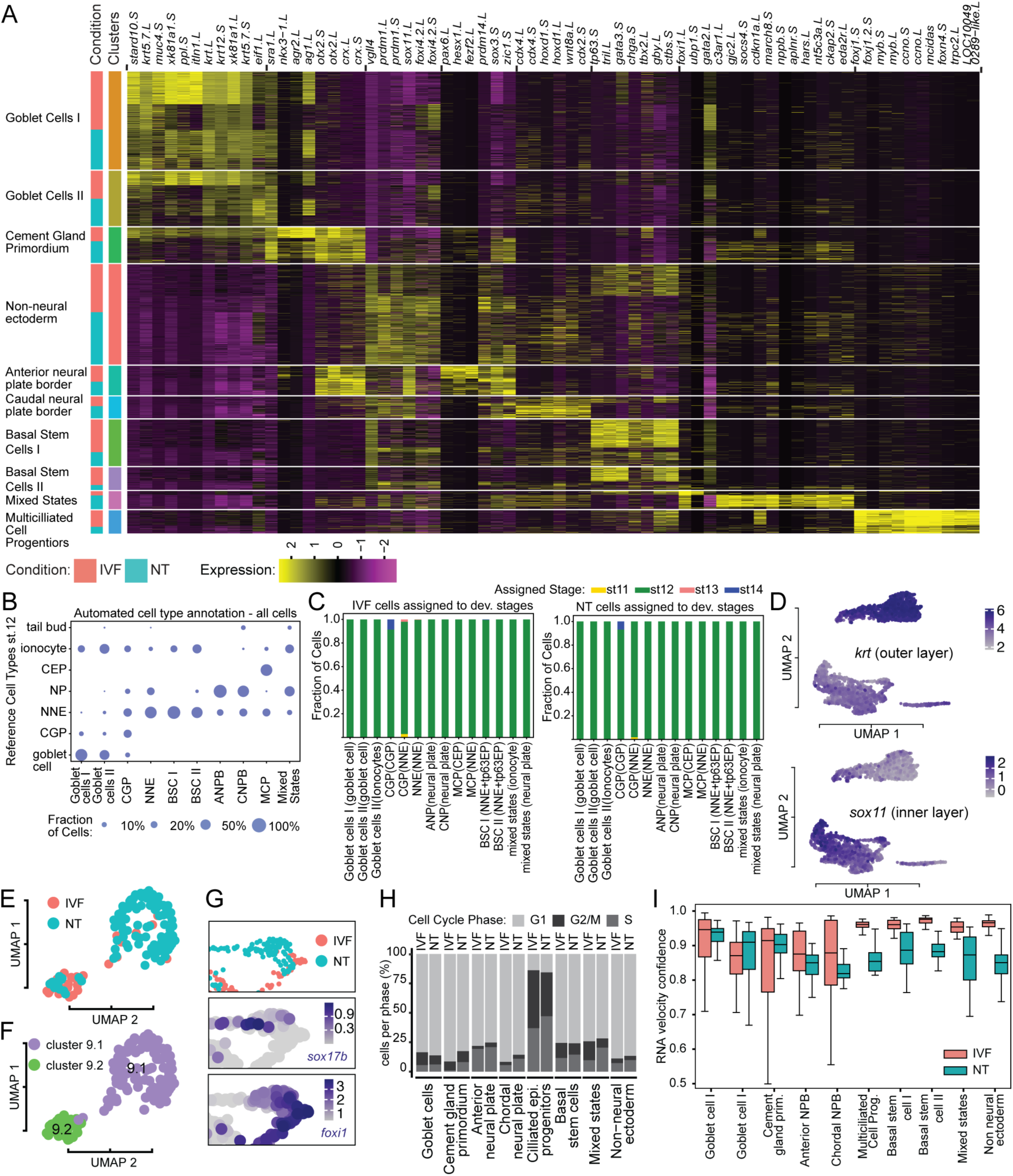
Differentiation defects vary across epidermal cell types and are associated with incomplete transcriptome reprogramming. **(A)** Heatmap showing scaled expression level of selected marker genes (columns) of the identified cell clusters (rows), used for cell type annotation. Cells ordered by cluster assignment and then by condition (IVF or NT). **(B)** Dotplot showing the fraction of cells from each cluster in our data (indicated on the x-axis) allocated to various cell types in the reference data (y-axis). Basal stem cells are missing from the X. tropicalis atlas, and they were mapped to non-neural ectoderm, which is the transcriptionally closest cell type at stage 12. **(C)** Fraction of cells from each cluster in our data (x-axis) mapping to the corresponding cell type in the X. tropicalis atlas at different stages, indicated by colors, for IVF (left) and NT (right) embryos. **(D)** Feature plots of krt and sox11 expression levels. (E) UMAP plot cluster 9. Coloured by condition IVF and NT. **(F)** UMAP plot cluster 9. Coloured by identified subclusters. **(G)** NT- and IVF- cells of Cluster 9 of panel D. **(H)** Cell-cycle analysis of NT and IVF epidermal cells per cluster using cyclone. Each bar represents proportion of cells in G1, G2/M and S-phase of the cell cycle. (I) Box plot of the RNA velocity confidence values per cell cluster, for IVF- and NT-cells separately. dev.=developmental; CEP=ciliated epidermal progenitor; NP=neural plate; NNE=non neural ectoderm; CGP= cement gland primordium; BSC=basal stem cell; ANPB=anterior neural plate border; CNPB= caudal neural plate border; MCP=multililiated cell progenitor

